# S-acylation targets ORAI1 channels to lipid rafts for efficient Ca2+ signaling by T cell receptors at the immune synapse

**DOI:** 10.1101/2021.02.03.429577

**Authors:** Amado Carreras-Sureda, Laurence Abrami, Ji-Hee Kim, Maud Frieden, Monica Didier, F. Gisou Van der Goot, Nicolas Demaurex

## Abstract

Efficient immune responses require Ca2+ fluxes across ORAI1 channels during engagement of T cell receptors (TCR) at the immune synapse (IS) between T cells and antigen presenting cells. Here, we show that ZDHHC20-mediated S-acylation of the ORAI1 channel at residue Cys143 is required for TCR assembly and signaling at the IS. Cys143 mutations reduced ORAI1 currents and store-operated Ca2+ entry in HEK-293 cells and nearly abrogated long-lasting Ca2+ elevations, NFATC1 translocation, and IL-2 secretion evoked by TCR engagement in Jurkat T cells. The acylation-deficient channel had reduced mobility in lipids, accumulated in cholesterol-poor domains, formed tiny clusters, failed to reach the IS and unexpectedly also prevented TCR recruitment to the IS. Our results establish S-acylation as a critical regulator of ORAI1 channel assembly and function at the IS and reveal that local Ca2+ fluxes are required for TCR recruitment to the synapse.

## Introduction

The development of an efficient immune responses by T lymphocytes require long-lasting Ca^2+^ elevations mediated by the plasma membrane (PM) channel ORAI1 during engagement of T cell receptors (TCR) at the immune synapse (IS) forming between T cells and antigen-presenting cells. Following TCR engagement, the Ca^2+^ depletion of the endoplasmic reticulum (ER) causes the ER-bound Ca^2+^ sensors STIM1-2 to oligomerize and to accumulate in ER-PM junctions, where they trap and gate the Ca^2+^-release-activated (CRAC) ORAI1 channel. The ensuing Ca^2+^ influx sustains long-lasting Ca^2+^ signals that initiate gene expression programs of T cell proliferation and differentiation. Proper ORAI1 function is essential for immunity in humans and patients with ORAI1 mutations suffer from severe combined immunodeficiency ^1^. Recent studies have revealed the structural rearrangements occurring within ORAI1 as STIM1 binding opens the channel pore and increases its selectivity for Ca^2+^ ^2, 3^, reviewed in ^4^. Crystal structure from the highly homologous *Drosophila* Orai1 channel revealed a hexamer of four concentric TM subunits, with pore-lining TM1 helixes bearing an acidic selectivity filter followed by hydrophobic and basic regions ^5, 6^. The closed structure is stabilized by multiple interactions between interlocking TM2 and TM3 helixes and peripheral TM4 helixes, bent in three crossed helical pairs extending in the cytosol. STIM1 binds to the external M4 helix, generating a gating signal transmitted by the TM2/TM3 ring to TM1, opening the channel pore and increasing its Ca^2+^ selectivity. The reversible switch of ORAI1 between a quiescent to an active state is highly regulated to avoid inappropriate Ca^2+^ fluxes at the wrong time or place (reviewed in ^4^).

Protein S-acylation, the reversible thioester linkage of a medium length fatty acid, often palmitic acid, on intracellular cysteine residues, dynamically controls the trafficking and gating of more than 50 ion channels by increasing the hydrophobicity of protein domains ^7^. S-acylation regulates ligand-gated (AMPA, GABA, Kainate, nAChR, NMDA, K_ATP_, P2×7 receptors), voltage-gated (Ca_V_, K_V_, K_Ca_ Na_V_), epithelial (ENaC), and water channels (AQP4). The S-acylation reaction is mediated by zinc-finger and DHHC-domain containing Protein AcylTransferases (PATs) at the ER and Golgi ^8^ and reversed by acyl protein thioesterases at the PM ^9^, with 23 PATs and 5 thioesterases isoforms identified in human so far ^10^. Due to the hydrophobic nature of the attached acyl moieties, protein S-acylation impacts the distribution of proteins in membrane microdomains and between intracellular membranes ^11^. Indirect evidences suggest that ORAI1 activity might also be controlled by S-acylation. First, the human isoform ORAI1 was identified by acyl-biotinyl exchange chemistry coupled to mass spectrometry as a robustly S-acylated proteins in primary human T cells ^12^ and human platelets ^13^. Second, the mouse Orai1 orthologue is reportedly to be S-acylated in neural stem cells and in a T-cell hybridoma ^14, 15^ according to the protein S-acylation database SwissPalm (https://swisspalm.org). Orai isoforms have two conserved Cys residues at potential S-acylation sites: Cys^143^, located in a privileged S-acylation position at the edge of the second transmembrane domain (TM2), and Cys^126^ within TM2. A third Cys residue, Cys^195^, sensitive to oxidation ^16, 17^, is exposed to the extracellular side and thus unlikely to be S-acylated. Among these three Cys residue, Cys^143^ is the only one conserved in C. elegans (Fig. S1).

Here, we show that the pore-forming subunit of the CRAC channel ORAI1 can undergo S-acylation at Cys^143^ and that this modification is required for efficient channel activity and for proper assembly of TCR at the immune synapse. Cys^143^ but not Cys^126^ substitutions prevents ORAI1 S-acylation mediated by PAT20. S-acylation-deficient ORAI1-C143A resides in cholesterol-poor membrane domains, forms tiny PM clusters upon store depletion, mediates reduced SOCE and I_CRAC_, and fails to reach the immune synapse in Jurkat T cells, severely impairing TRC assembly and synapse formation. S-acylation of ORAI1 therefore controls the recruitment and function of channels and receptors at the immune synapse to mediate efficient T cell responses.

## Results

### The ORAI1 channel can undergo S-acylation on Cysteine 143

Orai1 can potentially be S-acylated according to the SwissPalm 2.0 S-acylation database (https://swisspalm.org/) that compiles palmitoyl-proteomes ^18^. To validate that ORAI1 can undergo S-acylation, we assessed whether PEG-5k or tritiated palmitate could be incorporated by palmitoyl-thioester bonds into endogenous ORAI1 channels. HeLa cells were lysed in the presence of N-ethylmaleimide (NEM) to block free thiols, treated or not with hydroxylamine (HA) to break acyl-thioester bonds, and then with PEG-5k to label S-acylation sites. A mobility shift was observed in the presence of PEG-5k on western blots with anti-ORAI1 antibodies (Fig. 1A). Tritiated palmitate was detected by autoradiography in HeLa cells labelled for 2 h with ^3^H-palmitic acid and immunoprecipitated with anti-ORAI1 antibodies (Fig. 1B). These data show that endogenous ORAI1 channels incorporate palmitic acid and can be labelled by acyl exchange of the acyl moiety with PEG-5k. A single band of higher molecular weight was observed in the acyl-PEG assay, indicating that a single residue of the ORAI1 channel can be S-acylated in these conditions. The fact that a non-shifted ORAI1 band remains indicates that only a sub-population undergoes S-acylation under our experimental conditions.

**Fig. 1.**
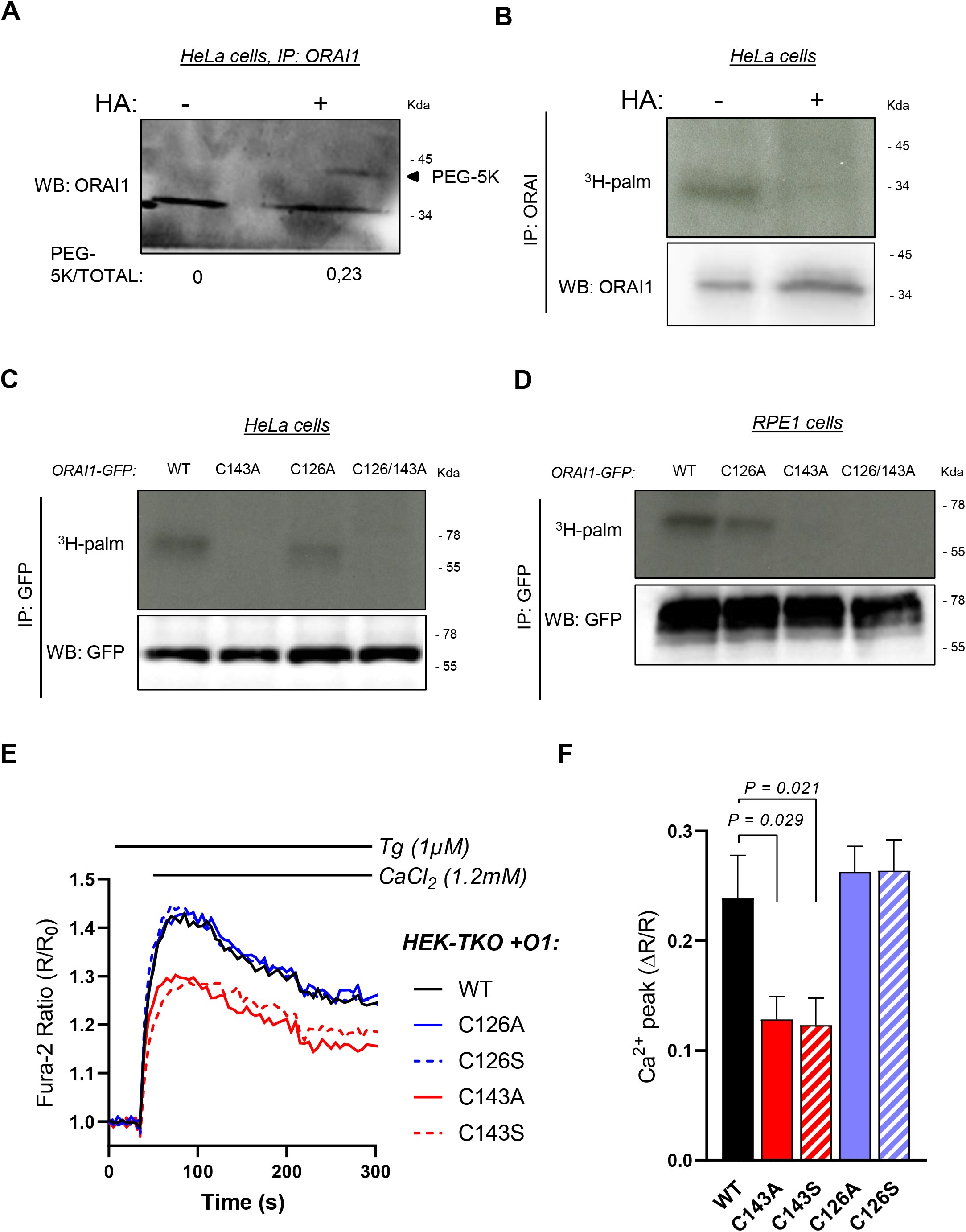
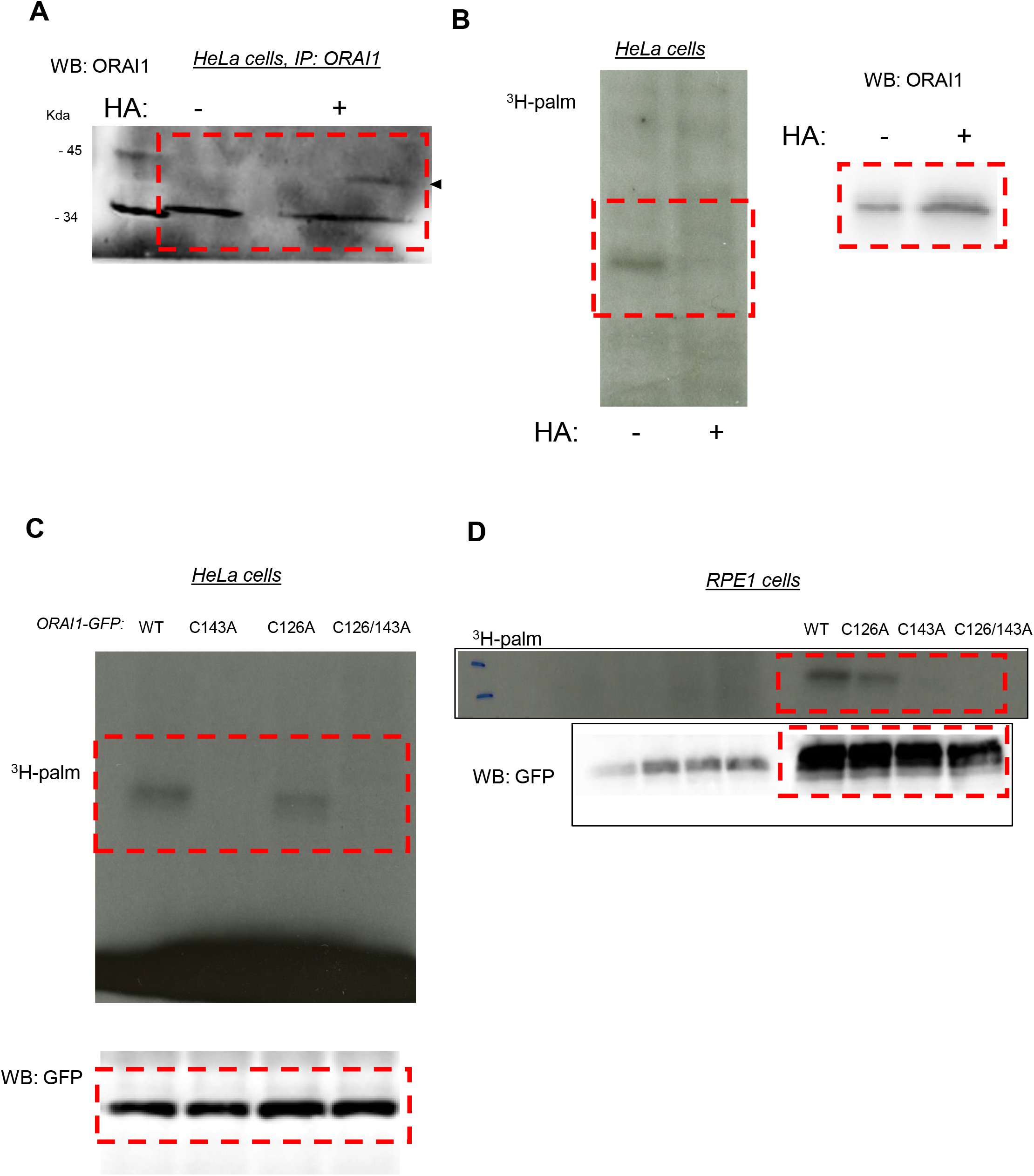
ORAI1 is S-acylated at Cysteine C143. (A) ORAI1 immunoblot of HeLa cells treated with PEG-5k to label S-acylation sites after exposure to NEM to block free thiols and then to hydroxylamine (HA) to break acyl-thioester bonds. (B) Western blot and corresponding autoradiogram of HeLa cells labelled for 2 h with ^3^H-palmitic acid with or without HA and immunoprecipitated with anti-ORAI1. (C, D) Western blots and corresponding autoradiograms of HeLa (C) and RPE-1 (D) cells expressing the indicated GFP-tagged ORAI1 mutants labelled with ^3^H-palmitic acid and immunoprecipitated with anti-GFP. Blots are representative of 3 independent experiments. (E) Normalized mean fura-2 responses evoked by Ca^2+^ readmission in HEK-TKO cells transiently transfected with the indicated ORAI1-GFP constructs and exposed to Tg. (F) Peak amplitude of the responses in E after background subtraction. Data are mean±SEM of 8 independent experiments. One way ANOVA Dunnett’s multiple comparisons test.

S-Acylation occurs on cysteine residues, present in ORAI1 at positions 126, 143 and 195, with C143 conserved up to *C*.*elegans* (Fig. S1A) and C195 facing the extracellular side (Fig. S1B). To test whether C126 and/or C143 are S-acylation sites we overexpressed ORAI1-GFP fusion proteins bearing substitutions at these residues in HeLa cells and evaluated ^3^H-palmitate incorporation by autoradiography. Cells expressing ORAI1-GFP bearing the C143A substitution or the double C126A/C143A mutation, but not the single C126A mutation, failed to incorporate ^3^H-palmitate (Fig. 1C). Identical results were obtained with these ORAI1-GFP mutants expressed in RPE1 cells (Fig. 1D), establishing that ORAI1 channels can undergo palmitoylation at residue C143.

### S-acylation potentiates ORAI1 channel function

S-acylation can alter ion channel trafficking, gating, and distribution in membrane lipids. To understand if S-acylation could affect ORAI1 activity we measured Ca^2+^ fluxes carried by ORAI1-GFP fusion constructs bearing substitutions at C126 and C143. In HEK-293 cells lacking all three ORAI isoforms (HEK-TKO, kindly provided by Rajesh Bhardwaj ^17^), expression of wild-type ORAI1 reconstituted Ca^2+^ fluxes upon store depletion (Fig. 1E-F). C143 substitutions by alanine or serine, but not C126 substitutions, reduced ORAI1-mediated SOCE (Fig. 1E-F). These findings were confirmed by alanine substitutions in HEK-293 cells stably expressing mCherry-STIM1 (mCh-STIM1) and ORAI1-GFP, (HEK-S1/O1). Although these cell lines were sorted for the same fluorescence and presented comparable STIM1 and ORAI1 levels as judged by epifluorescence microscopy (Fig. S1C), SOCE responses were strongly reduced in cells bearing the C143A mutation (Fig. 2A-B and S1D). We then recorded I_CRAC_ currents in HEK-S1/O1-WT and -C143A cell lines and observed a current density reduction of 5-fold in cells expressing the C143A mutant (Fig. 2C-D). The currents retained the inward rectification, positive reversal potential, and Gd^3+^-sensitivity characteristic of highly Ca^2+^ selective CRAC currents (Fig. 2C and 2E) but activated more slowly and failed to inactivate in a significant fraction of C143A cells (Fig 2F and S2). Our Ca^2+^ imaging and electrophysiological data thus establish that replacing the S-acylated Cys 143 residue within ORAI1 reduces the CRAC channel function.

**Fig. 2.**
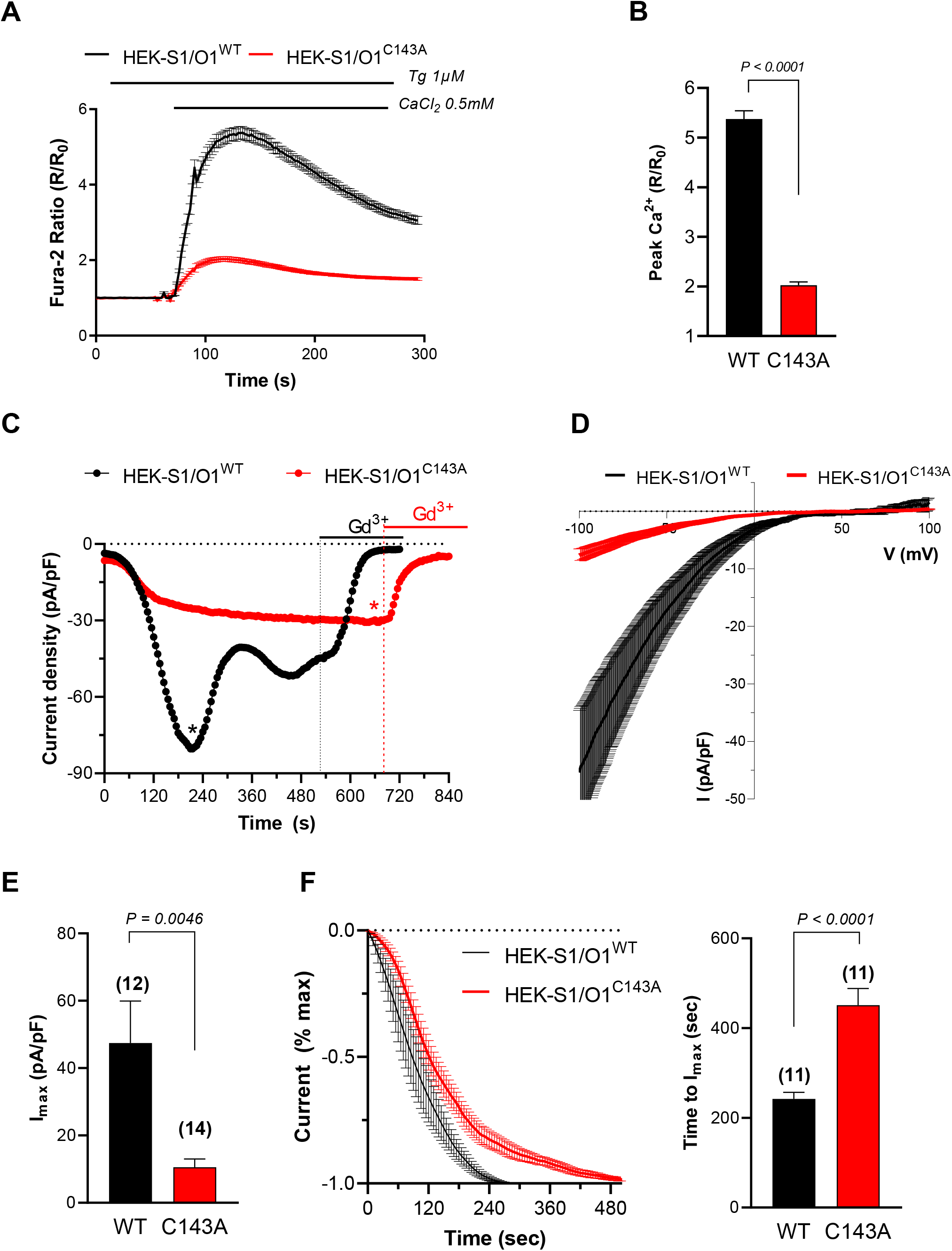
Preventing ORAI1 S-acylation reduces I_CRAC_ currents. (A) Normalized fura-2 responses evoked by Ca^2+^ readmission to Tg-treated HEK-293 cells stably expressing mCh-STIM1 and ORAI1-GFP (O1/S1) bearing or not the C143A mutation. (B) Peak amplitude of the responses in A. Data are mean±SEM of 196 (WT) and 198 (C143A) cells from 5 independent experiments. (C) Representative I_CRAC_ recordings of WT and C143A O1/S1 cells, measured every 5 seconds at −100 mV. I_CRAC_ was activated by cell dialysis with 10 mM BAPTA and blocked by 10 µM Gd^3+^. (D) Current-voltage relationship of the peak current in the cells shown in C (mean±SEM). (E) Peak current densities (I_max_) of WT and C143A O1/S1 cells after subtraction of basal or Gd^3+^-insensitive currents. (F) Time-course of current activation in cells without pre-activated currents. Left: Recordings were aligned to the first inflexion point and basal and maximal values set to 0 and 1, respectively. Right: Statistical evaluation of the activation time. Data are mean±SEM, number of cells is indicated on the graphs. Two-tailed unpaired Student’s *t*-test.

### Acylation increases ORAI1 cluster size, PM mobility, and affinity for lipid rafts

To gain insight into the underlying mechanism, we then recorded the formation of STIM1 and ORAI1 clusters during store depletion by TIRF microscopy. ORAI1-C143A clusters were tinier and occupied a smaller fraction of the TIRF plane (Fig. 3A-B). In contrast, the morphometric parameters of mCh-STIM1 clusters were not altered (Fig. S3). Lipid incorporation into proteins changes their lipophilic preference, and potentially their membrane mobility. To assess ORAI1 mobility in the PM we used fluorescence recovery after photobleaching (FRAP) and measured the lateral diffusion of ORAI1-GFP in HEK-S1/O1-WT and -C143A cell lines. The C143A mutant had a significantly lower diffusion coefficient indicative of a reduced mobility in membrane lipids (Fig. 3C). The addition of an acyl chain to transmembrane proteins increases their hydrophobicity, which may promote their association with lipid microdomains. To study whether S-acylation impacts ORAI1 lipid partitioning, we generated giant plasma membrane vesicles (GPMV) from HEK-293 cells transiently expressing ORAI1-YFP and measured the lipid distribution of the WT and mutated channel using the lipid raft and non-raft markers cholera toxin B and DiD, respectively. ORAI1-C143A co-localized less extensively with the raft marker in GPMVs from store-replete cells (i.e. not treated with thapsigargin), indicating that the acylation-deficient mutant has a reduced preference for lipid rafts (Fig. 3D). These results indicate that the acylation-deficient ORAI1-C143A mutant accumulates in cholesterol-poor lipid domains, has reduced mobility in the PM and forms tiny clusters upon store depletion. Preventing S-acylation thus impairs ORAI1 mobility in membrane lipids and its ability to form clusters during SOCE.

**Fig. 3.**
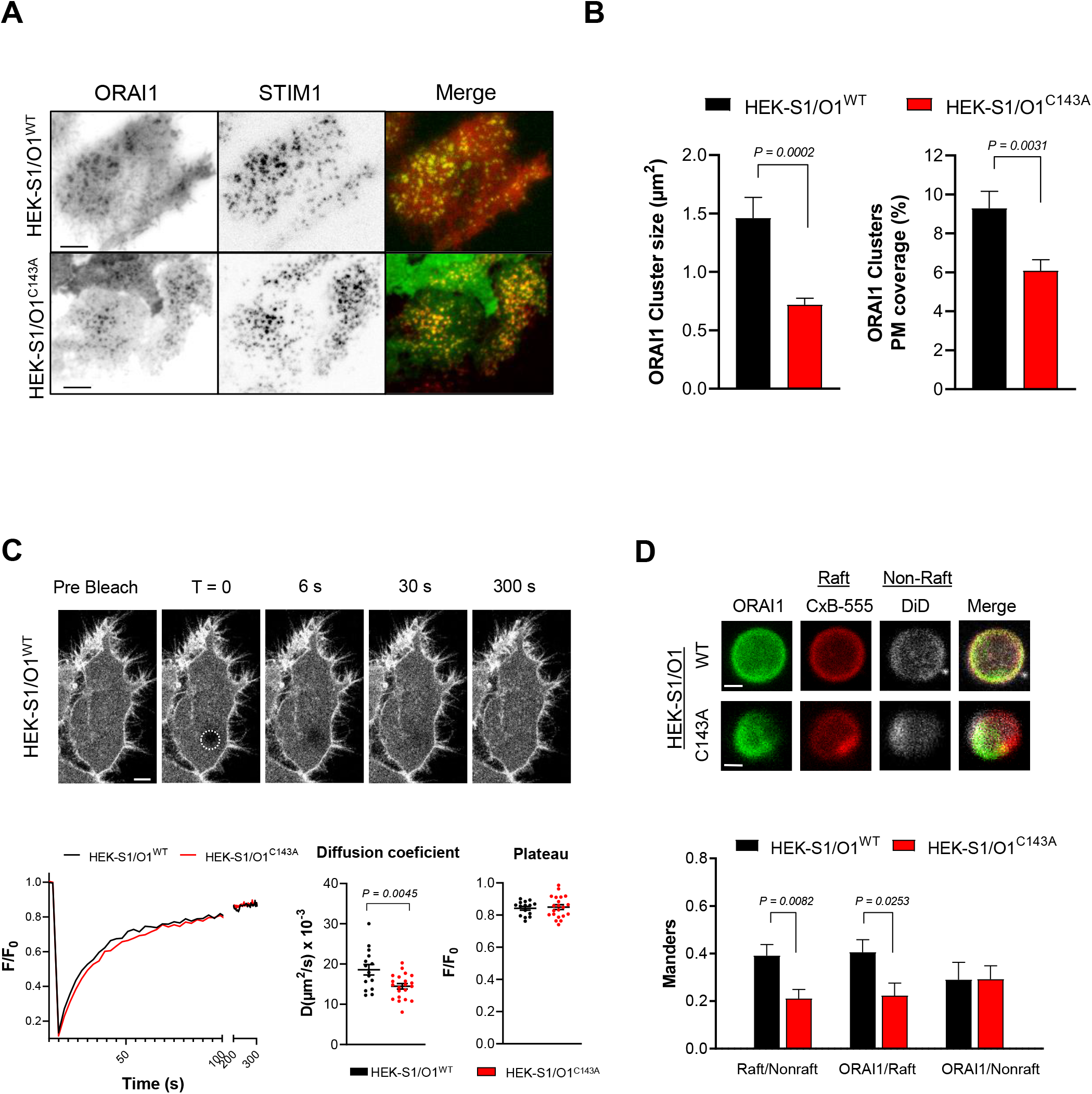
Preventing ORAI1 S-acylation reduces channel clustering and affinity for lipid rafts. (A) Representative TIRF images of WT and C143 O1/S1 cells exposed to 10 µM CPA for 10 min to induce mCh-STIM1 and ORAI1-GFP clustering Bars = 5 µm. (B) Averaged size of individual ORAI1-GFP clusters (left) and extent of PM covered by clusters (right) after CPA treatment. Data are mean±SEM of 29 (WT) and 30 (C143A) cells from 3 independent experiments. (C) FRAP recordings from WT and C143 O1/S1 cells. Top: representative GFP images. Bottom: representative fluorescence decay and recovery (left), diffusion coefficients (middle), and fluorescence plateau values (right). Data are mean±SEM of 21 (WT) and 15 (C143A) cells from 3 independent experiments Bars = 5 µm. (D) Lipid partitioning of ORAI1 in giant vesicles from HEK-293 cells transiently transfected with WT or C143 ORAI1-GFP. Top: representative fluorescence images of vesicles from cells expressing WT or C143 ORAI1-GFP (green) stained with cholera toxin subunit B (red) as raft marker and DiD (white) as non-raft marker top bar = 3 µm; bottom bar = 2µm. Bottom: Manders co-localization index for the indicated staining and conditions. Data are mean±SEM of 11 (WT) and 10 (C143A) vesicles from 3 independent experiments. Two-tailed unpaired Student’s *t*-test.

### Protein S-acyl transferase 20 (PAT20) mediates ORAI1 S-acylation

S-Acylation is exerted by DHHC-domain containing protein acyltransferases (PATs) proteins, which form a large family of enzymes containing 23 members. To identify the enzyme(s) promoting ORAI1 S-acylation, we transiently transfected ORAI1-GFP in HeLa cells stably expressing different PATs and measured palmitate incorporation in the immunoprecipitated channel by autoradiography. An enhanced palmitate incorporation was observed in cells expressing PAT3, PAT7 and PAT20 (Fig. 4A and S4A). Interestingly, a single band of ∼65 kDa was labelled in cells overexpressing PAT20, corresponding to full-length ORAI1 fused to GFP, while a ∼50 kDa band was predominantly detected in cells overexpressing PAT3 and PAT7, corresponding to a shorter form, likely Orai1β generated by alternative translation initiation at Met64 ^19^. We then assessed whether the enhanced ORAI1 S-acylation conferred by PATs overexpression could modulate channel activity. Enforced expression of PAT20 but not of PAT3 or PAT7 in HeLa cells enhanced SOCE (Fig. 4B) while siRNA against any of these PAT isoforms decreased SOCE equally (Fig. S4B). Importantly, PAT20 potentiated SOCE in HEK-293 cells stably expressing ORAI1-WT but not the C143A mutant (Fig. 4C), indicating that the gain of function conferred by increased PAT20-driven S-acylation requires this cysteine residue. PAT20-Myc immunoreactivity decorated reticular intracellular structures that co-localized with ORAI1-GFP clusters at the cell cortex (Fig. 4D), consistent with this enzyme mediating ORAI1 S-acylation. These experiments indicate that PAT20 mediates ORAI1 S-acylation and that this post-translational modification enhances SOCE.

**Fig. 4.**
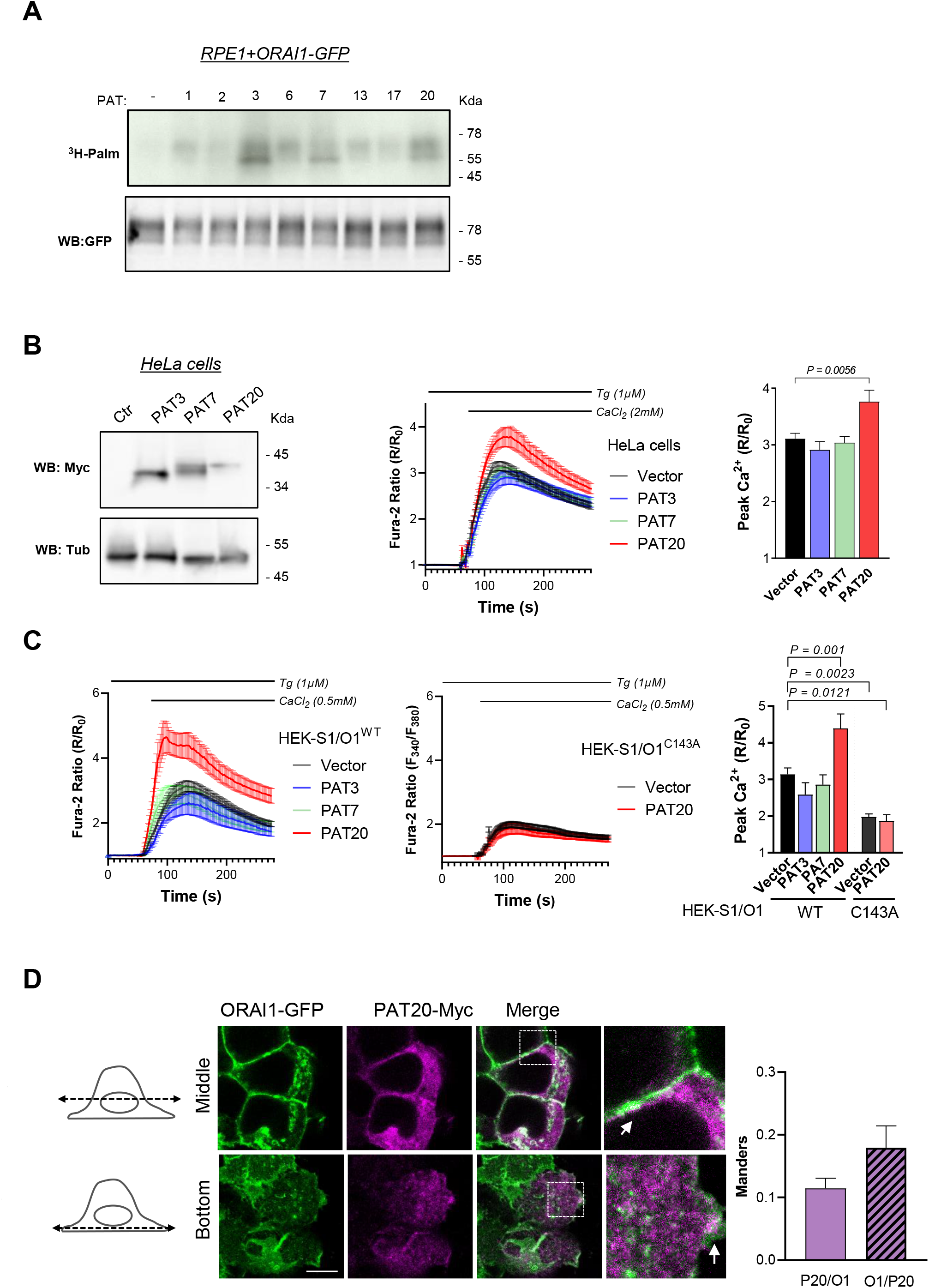
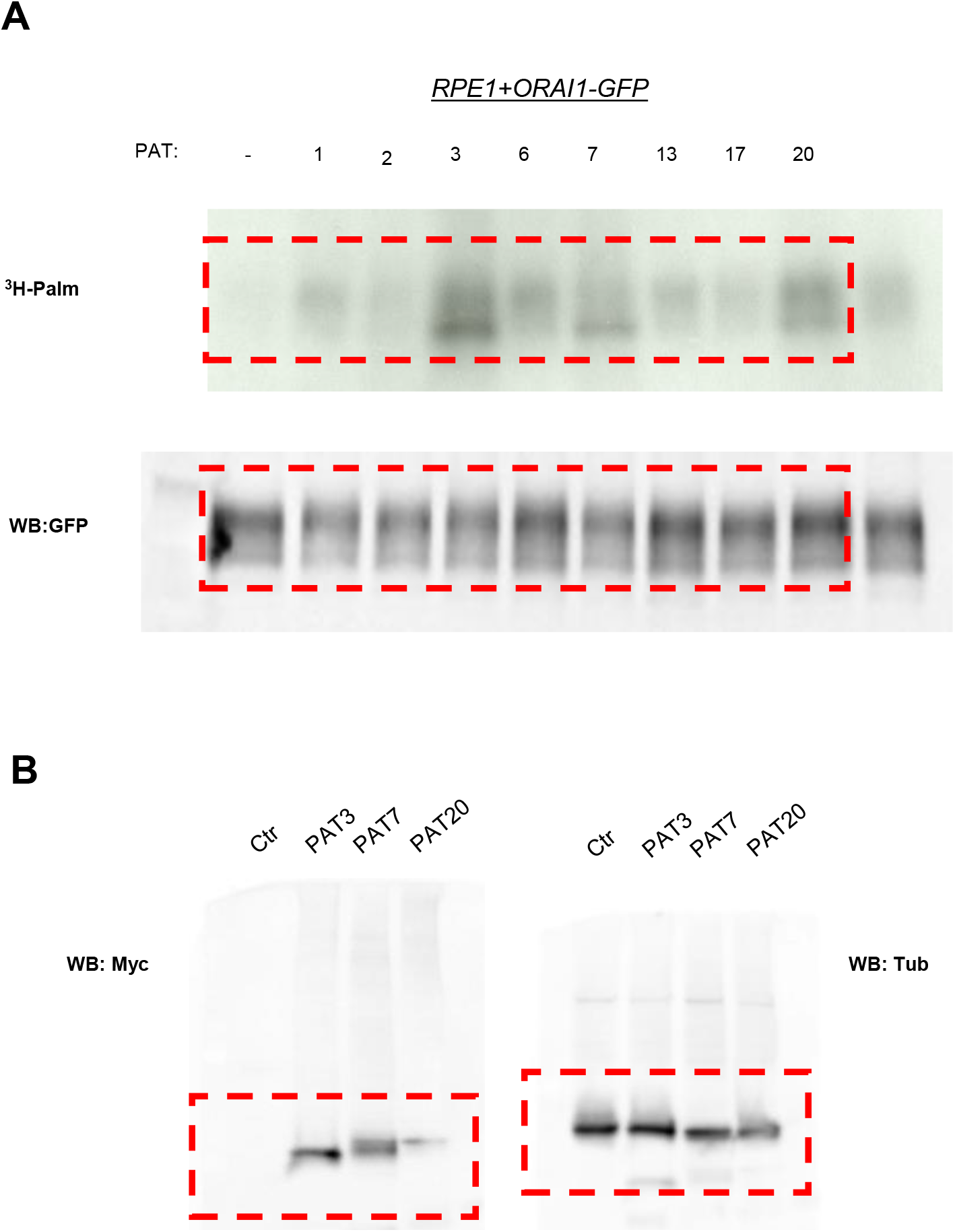
PAT20 S-acylates ORAI1 and modulates its activity. (A) Western blot and matching autoradiogram of RPE-1 cells expressing ORAI1-GFP plus the indicated PAT isoform, labelled with ^3^H-palmitic acid and immunoprecipitated with anti-GFP. Representative of 3 independent experiments. (B) Functional effect of PAT3, 7, and 20 expression. Western blot of HeLa cells expressing Myc-tagged PAT isoforms (left), averaged SOCE responses (middle), and peak amplitude (right). Data are mean±SEM of 49-74 cells from 3 independent experiments. (C) Averaged SOCE responses of WT (left) or C143A (middle) S1/O1 cells expressing these PAT isoforms and their peak amplitude (right). Data are mean±SEM of 31-129 cells from 5 independent experiments. (D) Confocal images of HeLa cells expressing Myc-tagged PAT20, treated or not with Tg (600s). Graphs show co-localization coefficients of Myc immunoreactivity with ORAI1-GFP, indicated by arrows on images. Bar = 10 µm. One way ANOVA Dunnett’s multiple comparisons test.

### ORAI1 S-acylation is required for TCR-mediated long-lasting Ca^2+^ elevations in Jurkat T cells

Orai1 activity is critical for the function of B and T lymphocytes, which fittingly express PAT20 but neither PAT3 nor PAT7 (Fig. S5A, http://www.humanproteomemap.org). To assess whether ORAI1 S-acylation by PAT20 impacts T cell function, we generated by CRISPR an ORAI1-deficient Jurkat T cell line, in which SOCE was severely blunted (Fig. 5A and S5B-C). Stable transduction of ORAI1-WT restored SOCE in these cells while ORAI1-C143A expressed at comparable levels was less effective (Fig. 5A and S5D). Activation of the T cell receptor (TCR) with CD3/CD28 beads evoked long-lasting Ca^2+^ elevations in CRISPR ORAI1 + ORAI1-WT stable cells (Fig. 5B). In contrast, cells reconstituted with ORAI1-C143A exhibited delayed responses of much smaller amplitude and duration upon TCR engagement. The Ca^2+^ signalling defect persisted when these cells were stimulated with TCR-coated beads in Ca^2+^-free medium, and subsequent Ca^2+^ readmission evoked minimal Ca^2+^ responses (Fig 5C). This indicates that the physiological Ca^2+^ signals engaged by the TCR receptors are severely affected in Jurkat cells expressing acylation-deficient ORAI1-C143A. Accordingly, PAT20 expression augmented SOCE in WT cells but had no effect in cells lacking ORAI1 (Fig. S5F). We then tested whether the downstream responses of T cells were similarly affected. Cells bearing the C143A mutation had reduced nuclear translocation of the transcription factor NFATC1 and IL-2 production following stimulation with Tg or CD3 beads (Fig. 5D-F-E and S5E). Furthermore, ORAI1 ablation prevented the potentiating effects of PAT20 expression on the IL-2 secretion evoked by Tg and CD3/CD28 beads (Fig. S5G). These results indicate that ORAI1 S-acylation at C143A is required for efficient activation of Jurkat T cells following TCR engagement.

**Fig. 5.**
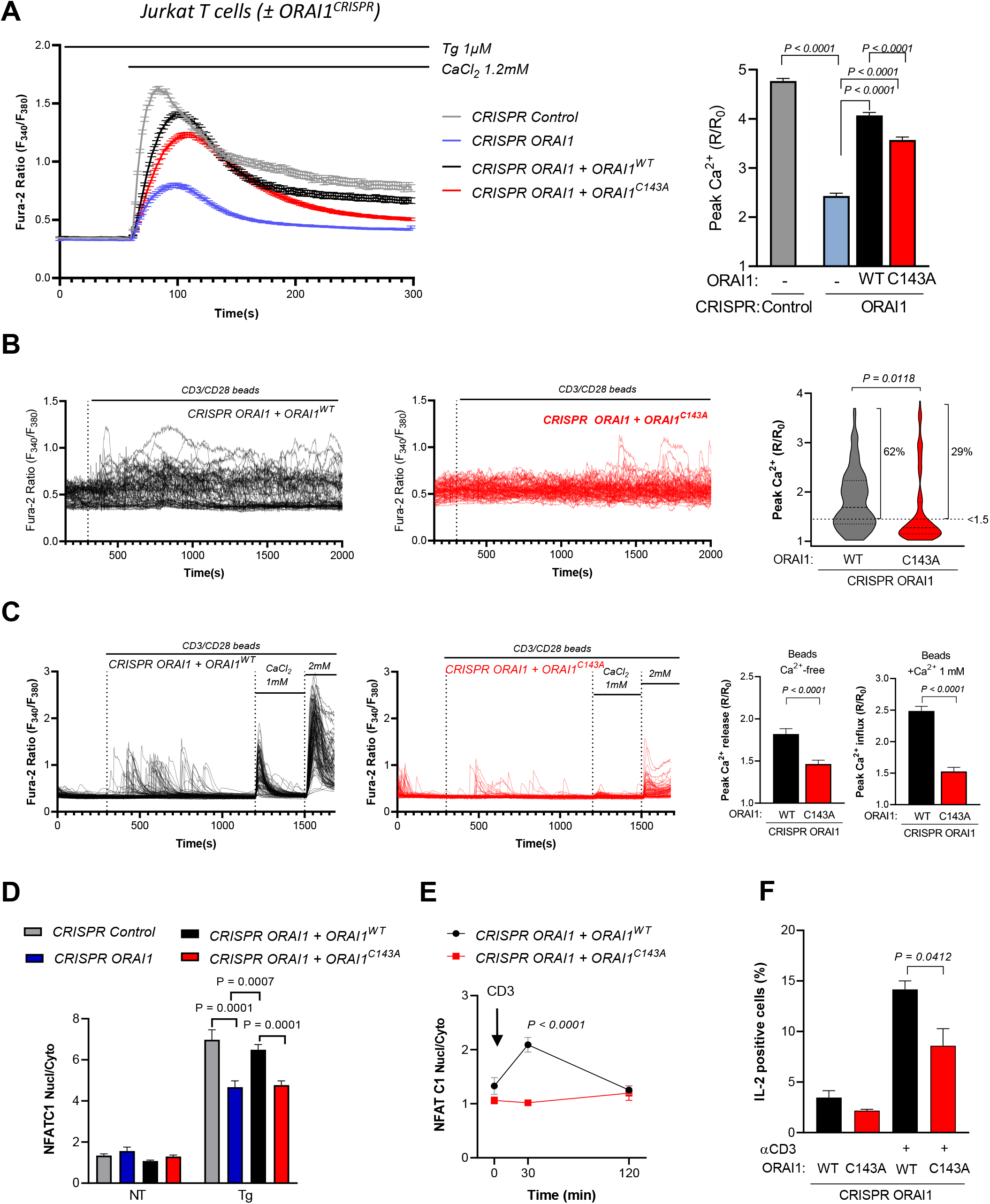
ORAI1 S-acylation promotes Jurkat T cell activation. (A) Averaged fura-2 responses and their peak amplitude evoked by Tg in Jurkat T cells lines generated by CRISPR with control or ORAI1-targeted guiding sequences and stably re-expressing either WT or C143A ORAI1-GFP. (B) Representative Fura-2 recordings of the indicated cell lines exposed to CD3/CD28-coated beads in physiological saline (left). Graph bars show the peak values evoked by CD3/CD28 beads in individual cells during the recording period. The percentages of cells with one or more elevation exceeding a threshold of 150% above basal is indicated. Data are from 102 cells (WT) and 124 cells (C143A) from 3 independent experiments. (C) Representative Fura-2 recordings of indicated cells exposed to CD3/CD28 beads in Ca^2+^-free media and then to 1 mM and 2 mM Ca^2+^ (left), and peak amplitude of these responses (right). Graph data are mean±SEM of 228 cells (WT) and 185 cells (C143A) from 3 independent experiments. (D) NFATC1 translocation evoked by Tg in the indicated cell lines. Data are mean±SEM of the nuclear to cytosol NFATC1-GFP intensity ratio of 58-161 cells from 4 independent experiments. (E) Time-course of NFATC1 translocation evoked by CD3 (OKT3 1µg/ml). Data are mean±SEM of 51-132 cells from 3 independent experiments. (F) IL-2 production evoked by CD3 (OKT3 1µg/ml). Data are mean±SEM of 3 independent experiments. One way ANOVA Dunnett’s multiple comparisons test (A and D) or two-tailed unpaired Student’s *t*-test.

### ORAI1 S-acylation sustains TCR assembly and signaling at the immune synapse

TCR activation triggers the formation of an immune synapse (IS) between T cells and antigen-presenting cells, a structure that maximizes the membrane contact area and organizes TCR and signalling proteins into concentric zones ^20^. ORAI1 channels are rapidly recruited into the IS ^21, 22^ and are required for the formation of dynamic actin structures ^23^ in a self-organizing process enabling long-lasting local Ca^2+^ signals to initiate gene expression programs that drive T cell proliferation ^24^. To test whether S-acylation impacts the recruitment of ORAI1 to the IS, we imaged CRISPR mediated ORAI1 deficient cells reconstituted with WT or mutant ORAI1-GFP during stimulation with antigen-coated beads or during plating on coverslips coated with anti-CD3 mAb ^25^. As previously reported, ORAI1-GFP accumulated at sites of bead contact, decorating dynamic cup structures labelled with SiR-Actin (Fig. 6A). Fewer SiR-Actin cups were observed in CRISPR ORAI1 Jurkat cells reconstituted with ORAI1-C143A, and the mutated GFP-tagged channel was not enriched at sites of contact when cups were detected (Fig. 6B). To better visualize the molecular organization of the IS, we performed TIRF imaging in coverslips coated with anti-CD3 mAb. ORAI1-GFP accumulated into contact zones surrounded by SiR-Actin rings. The formation of actin rings was severely compromised in cells reconstituted with the S-acylation-deficient ORAI1-C143A channel, which failed to accumulate at contact sites (Fig. 6C and S6A Suppl Video 1 and 2). The few rings forming in C143A mutant expressing cells had a comparable actin area (Fig. S6B) but contained less ORAI1-associated GFP fluorescence, detected predominantly in the centre of the IS (Fig. 6D and S6C). Unexpectedly, TCR cluster formation was also severely impaired in CRISPR ORAI1 cells reconstituted with acylation-deficient ORAI1. Although the two cell lines had comparable TCR surface expression (Fig. S6D), the number of CD3 immunoreactive dots within the IS was reduced 3-fold while their intra-IS distribution remained unaltered in cells reconstituted with ORAI1-C143A (Fig. 6E and S6C). The extent of co-localisation between ORAI-GFP and CD3 immunoreactivity was reduced in these cells, confirming the differential distribution of the two proteins in the IS (Fig. 6F and S6D). These data indicate that S-acylation is required for the recruitment of the ORAI1 channel to the IS and for the formation of TCR clusters that determine the intensity and duration of TCR signalling at the synapse (Fig. 6G).

**Fig. 6.**
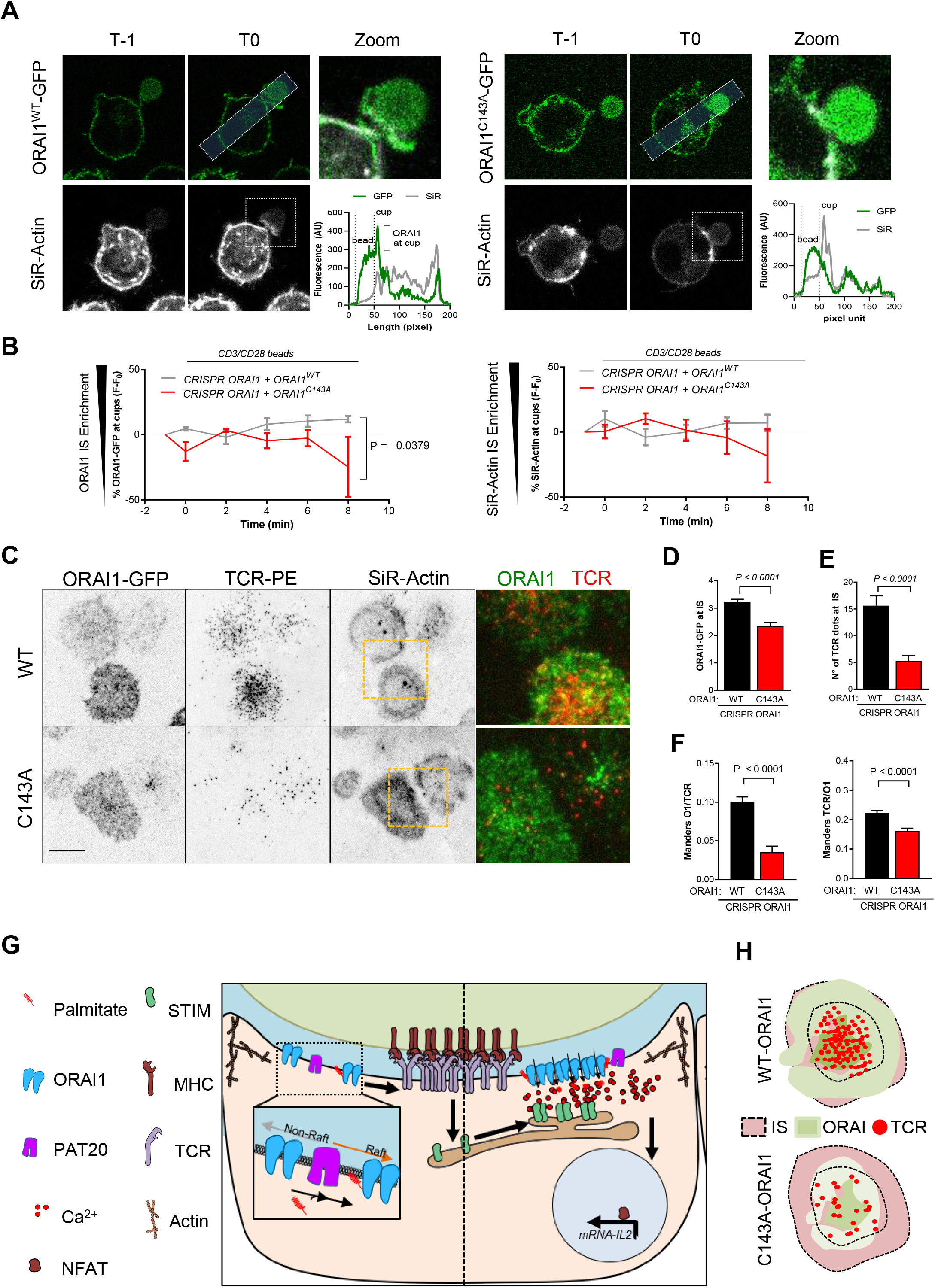
ORAI1 S-acylation regulates TCR clustering and signaling at the immune synapse. (A) Confocal images of ORAI1-deficient Jurkat T cells reconstituted with WT or mutant ORAI1-GFP, stained with SiR-Actin during initial contact with CD3/CD28-coated beads (visible by their autofluorescence in the GFP channel). Graphs show fluorescence intensities along IS-centred transcellular sections indicated by rectangles on GFP images. Dotted lines on SiR-Actin image indicate the zoomed regions. (B) Time-course of ORAI1-GFP (left) and Sir-Actin (right) accumulation at synapses forming in cells reconstituted with ORAI1-WT (5 cells) and ORAI1-C143A (3 cells). Two-ways ANOVA. (C) TIRF images of these cells stained with Sir-Actin and then with anti-TCR mAb after plating on activating coverslips coated with anti-CD3 mAb. Sketches show densities of TCR and ORAI1 in different concentric regions within the IS (Bar = 10 µm). (D) Averaged ORAI-GFP fluorescence and (E) numbers of TCR clusters within IS forming in these two Jurkat T cell lines. (F) Manders co-localisation index for TCR and ORAI1 WT or mutant. Data are mean±SEM of 28-61 cells from four independent experiments. Two-tailed unpaired Student’s *t*-test. (G) Scheme representing the effect of ORAI1 S-acylation on IS molecular composition and function. Addition of palmitate to ORAI1 channels by PAT20 targets the channel to lipid-ordered PM domains, promoting the formation of concentric ORAI1 and TCR clusters engaging MHC at the immune synapse. The resulting sustained local Ca^2+^ elevations (in red) induce nuclear translocation of NFATC1 to trigger IL-2 production. (H) Scheme representing the observed phenotype in TIRFF, where C143A mutant cells would accumulate less ORAI1 at the IS, it would redistribute in the centre and would have les TCR dots.

## Discussion

In this study, we show that S-acylation of ORAI1 at a single cysteine residue enhances the affinity of the channel for cholesterol rich lipid microdomains and promotes its trapping at the immune synapse, thereby enabling the local Ca^2+^ fluxes that control the proliferation of T cells. Using acyl-PEG exchange, palmitate incorporation, and mutagenesis, we show that ORAI1 can be chemically modified by S-acylation and identify the acylation site as Cys143 on the cytosolic rim of the second TM domain. Substitutions at Cys143 but not at Cys126 within TM2 prevented palmitate incorporation and decreased SOCE as well as I_CRAC_. A comparable inhibition was observed with cysteine-less ORAI1 in an earlier study focusing on Cys195 substitutions that prevent I_CRAC_ inhibition by hydrogen peroxide ^16^. These data indicate that the ORAI1 channel is S-acylated at Cys143 and that replacement of this residue, but not of the two other ORAI1 cysteines, prevents S-acylation and impacts channel function. Cys143 is the only cysteine conserved in all human isoforms and in ORAI1 homologs up to *C*.*elegans*, suggesting that S-acylation at this site is an evolutionary conserved function.

Using Ca^2+^ imaging and electrophysiology, we establish that ORAI1 S-acylation has a significant functional impact on the channel function. Substitutions at Cys143, but not at Cys126, decreased SOCE by 50% in HEK-293 cells when the channel was transiently expressed alone and by 80% when it was stably co-expressed with STIM1. SOCE was also reduced when the S-acylation-defective ORAI1-C143A was expressed in HEK-293 cells lacking all ORAI isoforms or in Jurkat T cells lacking ORAI1, firmly linking the SOCE defect to the ORAI1 Cys143 mutation. Patch-clamp recordings confirmed that I_CRAC_ currents were reduced by 80% by the mutation when ORAI1-GFP was stably expressed together with mCh-STIM1, at identical expression levels. ORAI1-C143A currents retained the characteristic inward rectification and high Ca^2+^ selectivity of CRAC channels but activated more slowly and failed to inactivate in a significant fraction of cells perfused with 10 mM BATPA in the pipette solution. This indicates that the C143A mutation does not grossly alter the gating or permeation properties of the CRAC channel. Instead, its main effect is to decrease the amplitude and to delay the activation of CRAC currents.

We further identify the zinc-finger and DHHC-containing S-acyltransferase zDHHC20 (PAT20) as mediating the S-acylation reaction. Among an array of PAT exogenously expressed in HeLa cells, PAT20 was the only isoform that increased the incorporation of tritiated palmitate into full-length ORAI1-GFP (Orai1α). PAT3 and PAT7 promoted palmitate incorporation in a lower band corresponding to a shorter form of ORAI1 (Orai1β). PAT20 co-localized with ORAI1-GFP at the cell cortex and unlike PAT3 and PAT 7 promoted SOCE when expressed. S-acylation by PAT20 thus positively modulates the activity of both endogenous and exogenously expressed ORAI1 channels. Importantly, SOCE potentiation was not observed when PAT20 was co-expressed with the acylation-deficient ORAI1-C143A mutant. This indicates that Cys143 is required for the potentiation by PAT20. Since ORAI1-C143A was co-expressed with STIM1 in these experiments, they also indicate that potential S-acylation sites on STIM1 are not relevant for the effect of PAT20. These data indicate that PAT20-mediated S-acylation at Cys143 enhances ORAI1 channel function.

Using biochemical and imaging approaches, we then show that mutating the Cys143 S-acylation site reduces the size of ORAI1 PM clusters during SOCE. We further show that the mutation reduces ORAI1 mobility in the PM and prevents its accumulation in ordered lipid domains rich in cholesterol. ORAI1 PM clusters are the macroscopic signature of ORAI1 trapping by STIM1, a dynamic event involving the entry and exit of ORAI1 particles into PM domains facing STIM1 molecules on apposed cortical ER cisternae ^26^. Molecularly, ORAI1 trapping reflects the interactions between the STIM1 CAD domain and ORAI1 C terminal tail, with residues within ORAI1 M4 helix being critical for trapping and gating. STIM1 clusters, on the other hand, reflect interactions between its lysin-rich C terminal tail and PM domains rich in negatively charged phospholipids such as PIP2. STIM1 therefore traps ORAI1 in PIP2-rich domains, while S-acylation increases ORAI1 affinity for cholesterol-rich domains. The increased mobility of S-acylated ORAI1 in cholesterol-rich domains likely increases its trapping by STIM1 into neighboring PIP2-rich domains since the escape probability of ORAI1 from STIM1-ORAI1 complexes is <1% ^26^. Increased ORAI1 retention at ER-PM contact sites would promote the formation of larger channel clusters and enhance transmembrane Ca^2+^ fluxes while preserving the biophysical properties of the channel, consistent with our observations. Alternatively, the formation of large clusters could reflect an increased affinity of S-acylated ORAI1 for STIM1 or increased lateral interactions between S-acylated channels leading to the formation of high-order channel multimers corresponding to the larger clusters.

By re-expressing the acylation-resistant ORAI1-C143A in ORAI1-deficient Jurkat T cells, we show that ORAI1 S-acylation is required for the efficient activation of T lymphocytes during TCR engagement. Replacing the single ORAI1 S-acylation site strongly reduced the long-lasting Ca^2+^ elevations driven by TCR engagement and the ensuing NFATC1 translocation and IL-2 production, signature markers of T cell activation. Unexpectedly, the responses were also reduced in Ca^2+^-free conditions, indicating that the inhibition of TCR signalling does not simply reflect the impaired channel function of ORAI1-C143A at the cell surface. Expressing PAT20 increased Ca^2+^ responses and TCR-induced IL-2 secretion in WT but not in ORAI1-deficient Jurkat T cells, confirming that ORAI1 S-acylation positively modulates TCR signalling. The reliance on S-acylation was most apparent at the IS, the specialized membrane contact area that form at the interface between T cells and an antigen presenting cells (APC). We observed three major synapse assembly defects in Jurkat T cells reconstituted with ORAI1-C143A. First, fewer synapses formed in ORAI1-C143A exposed to CD3-coated beads or plated on activating coverslips. Second, ORAI1-C143A was poorly recruited to the IS and the mutant channels accumulated in the IS centre. The IS contains a high percentage of highly ordered lipids ^27^ forming lipid rafts migrating to its periphery ^28^. S-acylation might target ORAI1 channels to these cholesterol-rich regions to optimize Ca^2+^ signalling efficiency at the synapse periphery (Fig. 6G). Third, the formation of TCR clusters was strongly reduced by the lack of ORAI1 S-acylation. This defect was unexpected as ORAI1 was not previously reported to control the molecular dynamics of TCR. During strong antigenic stimuli, TCR form clusters with associated scaffolding and signalling proteins that segregate in three concentric zones of the IS ^29, 30^. The clusters migrate from the periphery towards the centre of the IS where they are sorted for degradation ^31, 32^, the strength of signalling reflecting a balance between the formation of new clusters in the periphery and their disassembly in the centre ^33^. Defective ORAI1 targeting might impact TCR dynamics in several ways. In quiescent T cells, Ca^2+^ fluxes across ORAI1 channels might disrupt the CD3-lipid interactions that prevent spontaneous TCR phosphorylation ^34^, enhancing the activity state of TCR prior to their engagement. ORAI1 targeting to specialized PM domains such as filopodia might be required for this priming effect to occur. Alternatively, ORAI1 channels might control the rates of TCR recycling via endosomes by promoting the activity of Ca^2+^-dependent actin-severing proteins such as gelsolin. Our unexpected observation that ORAI1-C143A hinders TCR signalling even in the absence of extracellular Ca^2+^ suggests that ORAI1 might act from an intracellular location to promote endosomal recycling. The presence of ORAI1 channels in endosomes is well documented ^35^, but whether these channels mediate Ca^2+^ efflux from endosomes is not known. Preventing ORAI1 targeting could also impact the location or activity of integrin receptors such as ICAM-1, thereby indirectly altering the formation of TCR clusters. Further experiments are required to establish whether ORAI1 S-acylation promotes its endocytosis and whether S-acylation is dynamic or a one-off event that can impact the affinity of ORAI1 for accessory proteins or its potential interactions with other channels such as TRPC.

In summary, our findings reveal that the ORAI1 channel is regulated by S-acylation. The fatty acid addition is mediated by PAT20 and targets the channel to lipid-ordered PM domains rich in cholesterol, thereby facilitating channel trapping by STIM1 during cellular activation. The acylation-deficient channel failed to accumulate in the IS and prevented the formation of TCR clusters during TCR engagement, severely impeding the signals that drive T cell proliferation. We propose that S-acylation dynamically targets the ORAI1 channel to peripheral regions of the synapses rich in cholesterol to ensure efficient T cell signalling following TCR engagement.

## Materials and Methods

### Antibodies and reagents

The following reagents were used in this manuscript; Thapsigargin (T9033/CAY10522, Sigma); Ionomycin (I9657, Sigma); Phorbol 12-myristate 13-acetate (PMA) (79346, Sigma); Fura2-AM, (F1201, Invitrogen); Fluo-8, AM (21082, AAT Bioquest); SiR-Actin (Far Red, Spirochrome) Cyclopiazonic acid from Penicillium cyclopium, (c1530, Sigma); Gadolinium (G7532, Sigma); Vybrant® Alexa Fluor® 555 Lipid Raft Labeling Kit cholera toxin subunit B (V34404, Thermo-Fisher); Lipophilic Tracer Sampler DiD (L7781, Thermo-Fisher); Hoechst 33342 (H3570, Thermo-Fisher); Dynabeads^™^ Human T-Activator CD3/CD28 (11161D, Thermo-Fisher), GFP-trap agarose (GTA-10. Chromotek), - hydroxylamine (55460, Sigma, used at 0.5M), Zebra spin desalting columns (PIER89882, Pierce), 3H-palmitic acid (ART0129-25, American radio labelled chemicals), NEM (04559, Sigma), protein G (17-0618-01, GEHealthcare), 5kDa PEG (63187, Sigma). For protein detection either on Western blot or immunofluorescence, we used; NFATc1 (clone 7A6, MABS409, Sigma), TCR alpha/beta-PE (12-9986-42, eBioscience^™^), Anti-Cholera Toxin, B-Subunit (227040, Sigma), Myc-Tag (9B11) (2276, Cell-Signalling), gamma Tubulin (4D11) (MA1-850, Thermofisher), ANTI-FLAG® M2 (F1804, Sigma), anti-ORAI1 (600-401-DG9, rockland immunochemicals Inc), anti-GFP (SAB4301138, Sigma), anti-mouse-HRP and rabbit-HRP (1706516 and 172101, Bio-Rad (USA).

### Cell culture, cell lines and DNA constructs

Human embryonic kidney (HEK-293T) and Human retinal pigment ephitilial-1 (RPE1) cells were obtained from ATCC (CRL-11268, Manassas, VA, USA) maintained in Dulbecco’s modified Eagles medium (cat. no. 31966-021) supplemented with 10% fetal bovine serum and 1 % penicillin/streptomycin, and grown at 37°C and 5% CO_2_. HeLa cells purchased from the European collection of cell culture (ECACC) were grown in MEM Gibco (41090 in the same conditions. HEK-293 cells CRISPR triple knockout for ORAI1, 2 and 3 were a kind gift from Dr. Rajesh Bhardwaj, University of Bern. HEK-293T Stable cell lines expressing Cherry-Stim1 and hORAI1-WT or mutant C126A, C143A or C1267C143A were first infected with Cherry STIM1 p2K7 lentiviral vector, sorted, and then infected for the indicated mutants at a MOI of 2 and sorted for the same Cherry-STIM1 and ORAI1-GFP intensity. Indicated constructs were subcloned into a pWPT vector and co transfected with pCAG-VSVG/psPAX2 into HEK-293T cells to produce viral particles as described in ^36^. Briefly, indicated constructs were subcloned into a pWPT vector and co transfected with pCAG-VSVG/psPAX2 into HEK-293T cells to produce viral particles. After accumulation, ultracentrifugation and titration of the virus these were stored at −80°C. Jurkat T clone E6 cells were purchased from ECACC and grown in RPMI 1640 (21875-034 Life Technologies) supplemented with 10 FCS and 1% Pen/Strep. CRISPR Jurkat T cells were generated by stably expressing with lentiviral particles pLX-311-Cas9 construct (Addgene 96924) and transiently transfecting with Amaxa® Cell Line Nucleofector® Kit T (Ref: VCA-1002, Lonza) two sets of sgRNAs (Hs.Cas9.ORAI1.1.AA Ref: 224748421 / Hs.Cas9.ORAI1.1.AB Ref: 224748422, IDT). Single clone sorting and DNA sequencing were used to validate ORAI1 KO cells. ORAI1 rescue on CRISPR Jurkat T ORAI1 KO cells was performed by infecting at a MOI of 5 and FACS sorting for GFP fluorescence. To avoid clonal effects all cells used or generated in this study were pooled populations with the exception of HEK-TKO for ORAI1/2/3 or Jurkat CRISPR ORAI1 cells which were validated standard rescue, using either transient (HEK-293) or stable (Jurkat) expression. All cells sorted in this study were generated using a Beckman Coulter MoFlo Astrios integrated in PSL2 hood.

The ORAI1-yellow fluorescent protein (YFP) construct was purchased from Addgene (Cambridge, MA, USA; plasmid no. 19756). Site directed mutagenesis using the Pfu Turbo DNA polymerase from Agilent Technologies (Santa Clara, CA, USA; 600250) was used to introduce Cysteine mutants C143A, or S, and C126 A or s). Forward (fwd) and complementary reverse mutagenesis primers (Mycrosinth (Balgach, Switzerland) were as follows: C143A fwd: 5’-GCG CTC ATG ATC AGC ACC gcC ATC CTG CCC AAC ATC GAG GC-3’, C143S fwd: 5’-GCT CAT GAT CAG CAC CaG CAT CCT GCC CAA CAT CG-3’, C126A fwd: 5’-GCT CAT CGC CTT CAG TGC Cgc CAC CAC AGT GCT GGT GGC-3’, C126S fwd: 5’-GCT CAT CGC CTT CAG TGC CaG CAC CAC AGT GCT GGT GGC-3’. All plasmids encoding for human DHHC1, 2, 3, 6, 7, 13, 17 and 20 were Myc tagged in the N-terminus in pcDNA3 vectors, kindly provided by the Fukata lab.

### Radiolabeling 3H-palmitic acid incorporation

To follow S-acylation, transfected or non-transfected cells were incubated 1 hour in medium without serum (Glasgow minimal essential medium buffered with 10 mM Hepes, pH 7.4), followed by 2 hours at 37°C in IM with 200 µCI /ml 3H palmitic acid (9,10-3H(N)), washed with cold PBS prior immunoprecipitation overnight with anti-ORAI antibodies and protein G-beads or anti-GFP agarose-coupled beads. Beads were incubated 5 minutes at 90°C in reducing sample buffer prior to SDS-PAGE. Immunoprecipitates were split into two, run on 4-20% gels and analysed either by autoradiography (3H-palmitate) after fixation (25% isopropanol, 65% H2O, 10% acetic acid), gels were incubated 30 minutes in enhancer Amplify NAMP100, and dried; or Western blotting.

### Acyl-Peg-exchange

To block free cysteine, cells were lysed and incubated in 400 μl buffer (2.5% SDS, 100 mM HEPES, 1 mM EDTA, 40mM NEM pH 7.5, and protease inhibitor cocktail) for 4 h at 40°C. To remove excess unreacted NEM, proteins were acetone precipitated and resuspended in buffer (100 mM HEPES, 1 mM EDTA, 1% SDS, pH 7.5). Previously S-acylated cysteines were revealed by treatment with 250 mM hydroxylamine (NH_2_OH) for 1 hour at 37°C. Cell lysates were desalted using Zebra spin columns and incubated 1 hour at 37°C with 2mM 5kDa PEG: methoxypolyethylene glycol maleimide. Reaction was stopped by incubation in SDS sample buffer for 5 minutes at 95°C. Samples were separated by SDS-PAGE and analysed by immunoblotting.

### Ca^2+^ imaging and plate reader

Calcium assays in single cell live imaging were performed as described previously ^37^. Briefly, cells were loaded with 3 µM Fura-2-AM, in modified Ringer’s for 30 min at room temperature (RT). 340/380 nm excitation and 510 ± 40nm emission ratiometric imaging was performed every 2 seconds. SOCE activity was triggered by emptying the ER stores by blocking SERCA with Thapsigargin in a Ca^2+^-free solution containing 1 mM EGTA instead of 2 mM CaCl_2._Extracellular calcium addition revealed ORAI1 activity. Jurkat cells attachment to the coverslip was achieved by seeding 200.000 cells in 25mm poly-L lysine coated coverslips for 25 minutes at RT. When indicated, Jurkat cells were transfected with YFP cameleon (YC 3.6) calcium cytosolic probe to measure cytosolic calcium. YC 3.6 was excited at 440nm and emission was collected alternatively at 480 and 535 nm. Calcium imaging in HEK-293 cells Triple Knockout for ORAI1,2 and 3 was performed in plate reader using Fura2 as calcium Dye. Fluorescence was measured using a 96-well microplate reader with automated fluid additions at 37 °C (FlexStation 3, Molecular Devices).

### TIRF imaging

TIRF imaging to determine ORAI and STIM clusters in HEK-293 cells S1/O1 was performed on a Nikon Eclipse Ti microscope equipped with a Perfect Focus System (PFS III) using a 100× oil CFI Apochromat TIRF Objective (NA 1.49; Nikon Instruments Europe B.V.). To observe STIM1/ORAI1 clusters cells were bathed with CPA 10µM and imaged every 20 seconds in calcium free solution. Jurkat T cell lines were used in TIRF to image immune Synapse. Actin was imaged by loading Jurkat T cells (1 x 10^5^) with SiR-Actin (500nM) for 30 min at 37 °C 5% CO2. 500.000 Jurkat T cells were seeded on CD3 (OKT3, 1µg/ml) coated glass coverslips (25mm) at the beginning of the experiment and imaged every 1 minutes for 25 minutes. Then, TCRα/β-PE conjugated antibody was added (1:1000) in order to image TCR cluster formation in the IS in living cells. For both cell lines, ORAI1 was imaged using ZET488/10 excitation filter (Chroma Technology Corp.). STIM1 cherry (in HEK-293 cells) or TCR clusters were imaged using a ZET 561/10 excitation filter (Chroma Technology Corp., Bellows Falls, VT). SiR-Actin was measured using 640 nm laser line. All emission signals were collected by a cooled EMCCD camera (iXon Ultra 897, Andor Technology Ltd). All experiments were performed at room temperature (22–25°C).

### Confocal live imaging, FRAP and Beads assay

Confocal time lapse microscopy was used to image JurkaT cells with CD3/CD28 beads and Fluorescence recovery after photobleaching. For IS we used Jurkat ORAI1 expressing cells loaded with SiR to visulaize Actin (same protocol as for TIRF). 500.000 Jukrat cells were seeded on Poly-L lysine coated glass coverslips (25mm) for 25 minutes at RT. Beads (1:1 bead cell ratio) were added after 2 minutes of imaging of the experiment and imaged every 2 in a XYTZ configuration (Z stack spanning all cell with 1 µm of thicknes) for minutes for 25 minutes at RT. Images were obtained in a Nikon A1r Spectral with a 60x 1.4 CFI Plan Apo Lambda WD:0.13mm objective using 488 and 639 laser lines. Image analysis was performed by selecting in focus stacks where the bead was observed and measuring ORAI1 or SiR-Actin fluorescence over time in the bead contact area normalized to the fluorescence of the opposite pole as described previously ^38^.

FRAP was performed in HEK-293 S1/O1 under resting condition using the same microscope. ORAI1 FRAP was accomplished by following the protocol previously described ^39^. Briefly, we used a live chamber at 37°C and 5% CO2. Pinhole was settled at 1AU and images were sampled every 3 seconds for 100 images. Bleaching was for 20 seconds (488nm 100% output) after 1 minute of basal acquisition. ROI of interest was compared to the same size ROI in the same field of view and normalized to basal. Traces were fitted with an exponential one-phase association model to obtain the half-life, τ1/2 and fluorescence recovery. Diffusion coefficient was calculated with the formula D = 0.224r^2^/(τ1/2), in which r is the radius of the bleached circle region as described in ^39^.

### Flow cytometry

Cytometry calcium experiments on Jurkat cells were performed by incubation with Fluo8 (2µM 30 min, RT) and washed for 15 minutes in a calcium containing solution. BD Accuri C6 was used to measure calcium movements over time by setting the flow at 1µL per second. Every experiment started with 5 x 10^5^ cells in 510 µL of calcium free solution (1mM EGTA). After 1 min 50µL of Tg 10µM was added to empty ER stores. After 300 seconds we added 100 µL of CaCl_2_ (Final concentration 2.5mM) to reveal SOCE. IL-2 measurements were performed as described previously ^40^. Briefly, Jurkat cells (50.000) were seeded in pre coated CD3 (OKT3) round bottom 96 cell plates for 2 days. Cells were then fixed (PFA 4%) and perm/blocked with PBS-2%BSA 0.5% Saponin previous to IL-2 PE incubation. IL-2 FACS measurements were acquired in a BDLSR Fortessa unit.

### Giant Plasma Membrane Vesicles (GPMV)

GPMVs were formed and analysed following this protocol ^41^. Briefly, HEK-293 cells were seeded in poly-L-lysine coated 25mm glass coverslips and transfected with YFP-ORAI1 WT or the C143A mutant. The day after cells were washed with GPMV buffer (150mM NaCl, 2mM CaCl2 and 10mM HEPES, pH: 7.4) and incubated for 1h at 37°C 5% CO2 with a vesiculation buffer (25mM PFA, 2mM DTT,). Cell super natant was then spun for 30 minutes at 20.000 x g and incubated with Alexa-555 Cholera Toxin B subunit and DiD (Far red) lipid staining markers for 10 minutes on ice. Imaging was performed at 10°C using Open Perfusion Microincubator (PDMI-2, Medical Systems, Greenvale, NY) temperature controller to enhance lipid partitioning. Vesicles were imaged using a Nikon A1r Spectral with a 60×1.4 CFI Plan Apo Lambda WD:0.13mm objective using 488, 551 and 639 laser lines.

### Electrophysiology

I_CRAC_ currents were recorded using the whole-cell configuration in HEK-293 cells stably expressing mCherry-STIM1 and ORAI1-GFP (O1/S1) bearing or not the C143A mutation. The cells were trypsinized, seeded on 35 mm dishes (Corning, NY, USA) and incubated overnight at 37 °C to allow attachment of separated cells. The experiments were performed at room temperature. Pipettes were pulled from 1.5 mm thin-wall glass capillaries (GC150TF, Harvard Apparatus) using a vertical PC–10 Narishige puller to obtain a resistance between 2-4 MΩ. Currents were recorded with pCLAMP 10.7 software (Molecular Devices, Sunnyvale, CA, USA), using the Axopatch 200B amplifier (Axon Instruments, Molecular Devices) with a low–pass filtering at 1 kHz, and digitized with the Axon Digidata 1440A at 1 ms. Voltage ramps of 180 ms were applied from –120 to +100 mV every 5 seconds from a holding potential of 0 mV. Peak current densities (I_max_) were measured at –100 mV after subtraction of basal or 10 µM GdCl_3_-insensitive currents. The standard 10 mM Ca^2+^ recording solution contained 130 mM NaCl, 5 mM KCl, 1 mM MgCl_2_, 10 mM CaCl_2_ and 10 mM HEPES (300–310 mOsm, pH 7.4 adjusted with NaOH). The intracellular pipette solution contained 130 mM Cs methanesulfonate, 8 mM MgCl_2_, 10 mM BAPTA, and 10 mM HEPES (290–300 mOsm, pH 7.2 adjusted with CsOH).

### Immunofluorescence

PAT20 and ORAI1 immunofluorescence was performed in HeLa cells co-transfected with ORAI1-YFP and PAT20-myc. After 24h of transfection, cells were fixed (Pfa 4%) for 20 min at RT, then permeabilized (PBS-BSA0.5% + NP-40 0.1%) for 10 minutes and then blocked (PBS-BSA0.5% + FBS 5%) for 1h at RT. Then cells were incubated with primary antibodies O/N at 4°C then with secondary 1:1000 with Hoesch 1:5000 for 1h at RT. For NFAT translocation Jurkat cells were treated with the indicated compounds (Tg 1µM or CD3 plastic coated plates) for the indicated times and seeded into poly-L lysine coated coverslips for 15 minutes at RT. Immunofluorescences for NFATC1 were performed as described for HeLa cells. NFATC1 analysis was done by dividing the nuclear to the cytosolic (total-nuclear) pixel intensity per cell into 3 to 5 randomized fields per condition. Images were obtained in a LSM700 Nikon microscope.

### Image analysis and statistics

All images were analysed using ImageJ software. Co-localisation and particle concentric counting for TCR were performed by applying a previously described macro ^42^.

## Data availability

The data that support the findings of this study are available from the corresponding author upon reasonable request.

## Acknowledgements

We are grateful to Cyril Castelbou for the technical assistance, the bioimaging and flow cytometry facilities (Geneva Medical Centre). This work was funded by the Swiss National Foundation [grant number 310030_189042 (to ND) and SNF 310030B_176393 and 310030_192608 European Research Council under the European Union’s Seventh Framework Programme (FP/2007-2013) / ERC Grant Agreement n. 340260 - PalmERa’ (to GvG.).

## Author contributions

ACS, Conception and design, Acquisition of data, Analysis and interpretation of data, Drafting or revising the article; JK, MB, LA, & MF, Acquisition of data, Analysis and interpretation of data; GvG & ND, Conception and design, Analysis and interpretation of data, Drafting or revising the article.

## Figures Legends

**Supplementary Fig. 1 (related to Fig. 1).**
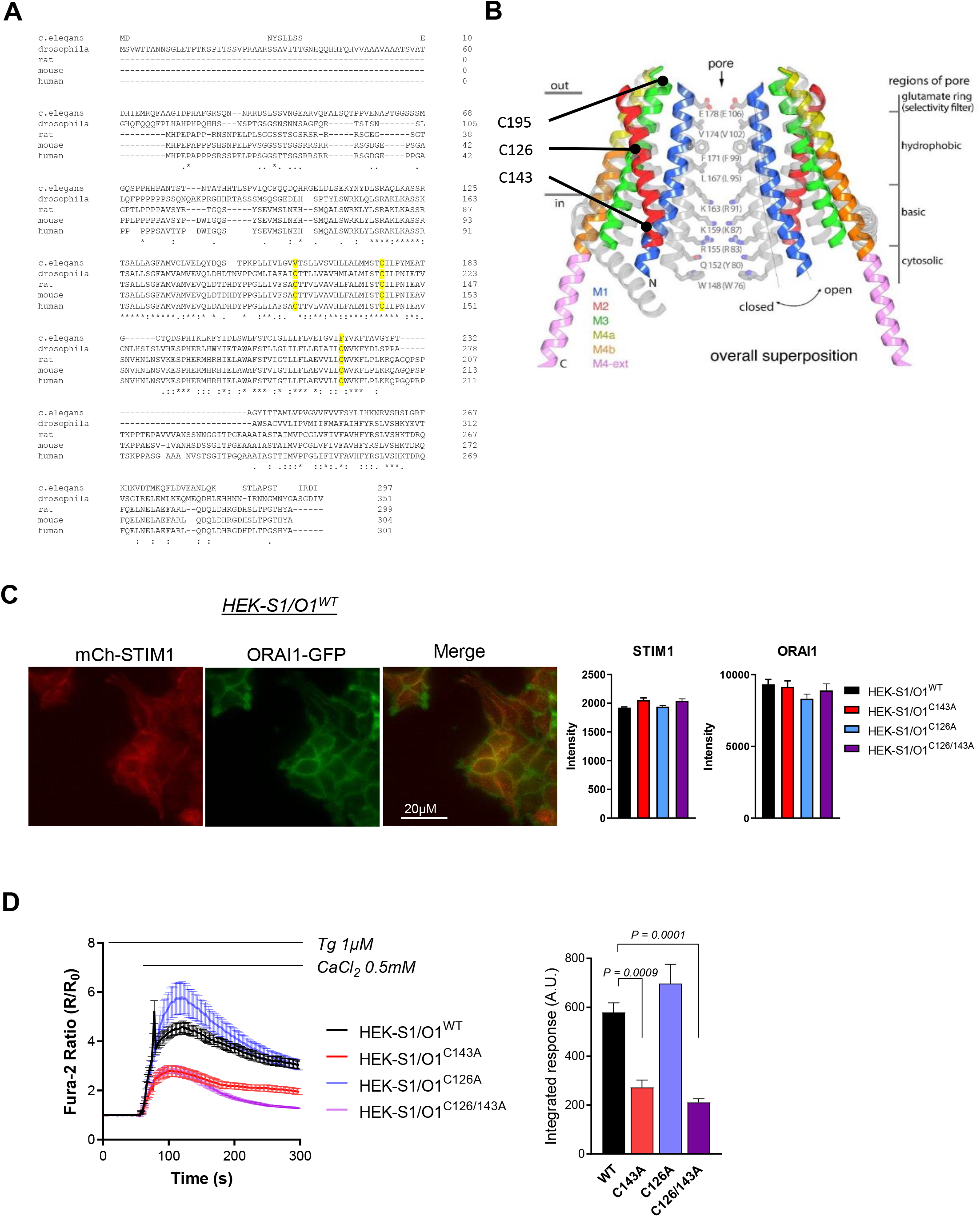
(A) Orai1 protein sequences aligned with CLustalW algorithm for the indicated organisms. Cysteines susceptible to be S-acylated are highlighted in yellow. (B) Schematic ORAI1 representation. Superimposed structures of the WT and H206A dOrai (PDB ID: 4HKR and 6BBF) conformations in ribbon representation highlighting cysteine residues at position 126, 143 and 195 (C) Fluorescence images of WT O1/S1 cells (left) and averaged mCh-STIM1 and ORAI1-GFP fluorescence of the different O1/S1 stable cell lines. (D) Averaged fura-2 responses (left) and peak amplitude (right) of O1/S1 cells bearing or not the indicated ORAI1 mutation(s). Data are mean±SEM of 44-82 cells from two independent experiments. One way ANOVA Dunnett’s multiple comparisons test.

**Supplementary Fig. 2 (related to Fig. 2).**
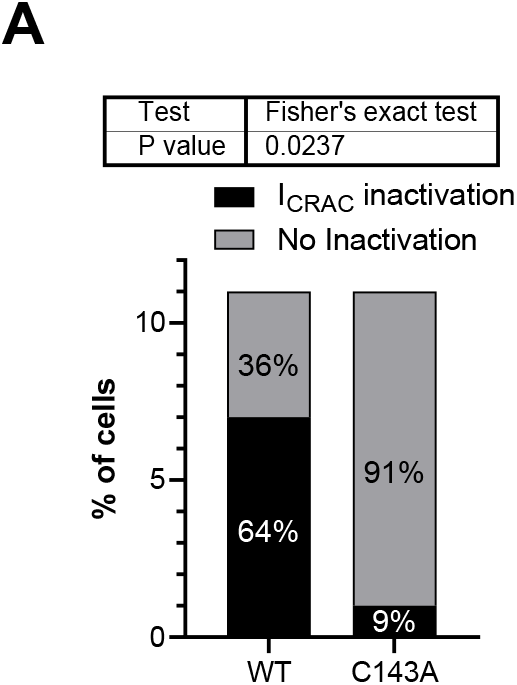
(A) Number of recorded WT and C143A O1/S1 cells exhibiting or not I_CRAC_ inactivation. Chi-square p value: 0.0237, two-sided Fisher’s exact test.

**Supplementary Fig. 3 (related to Fig. 3).**
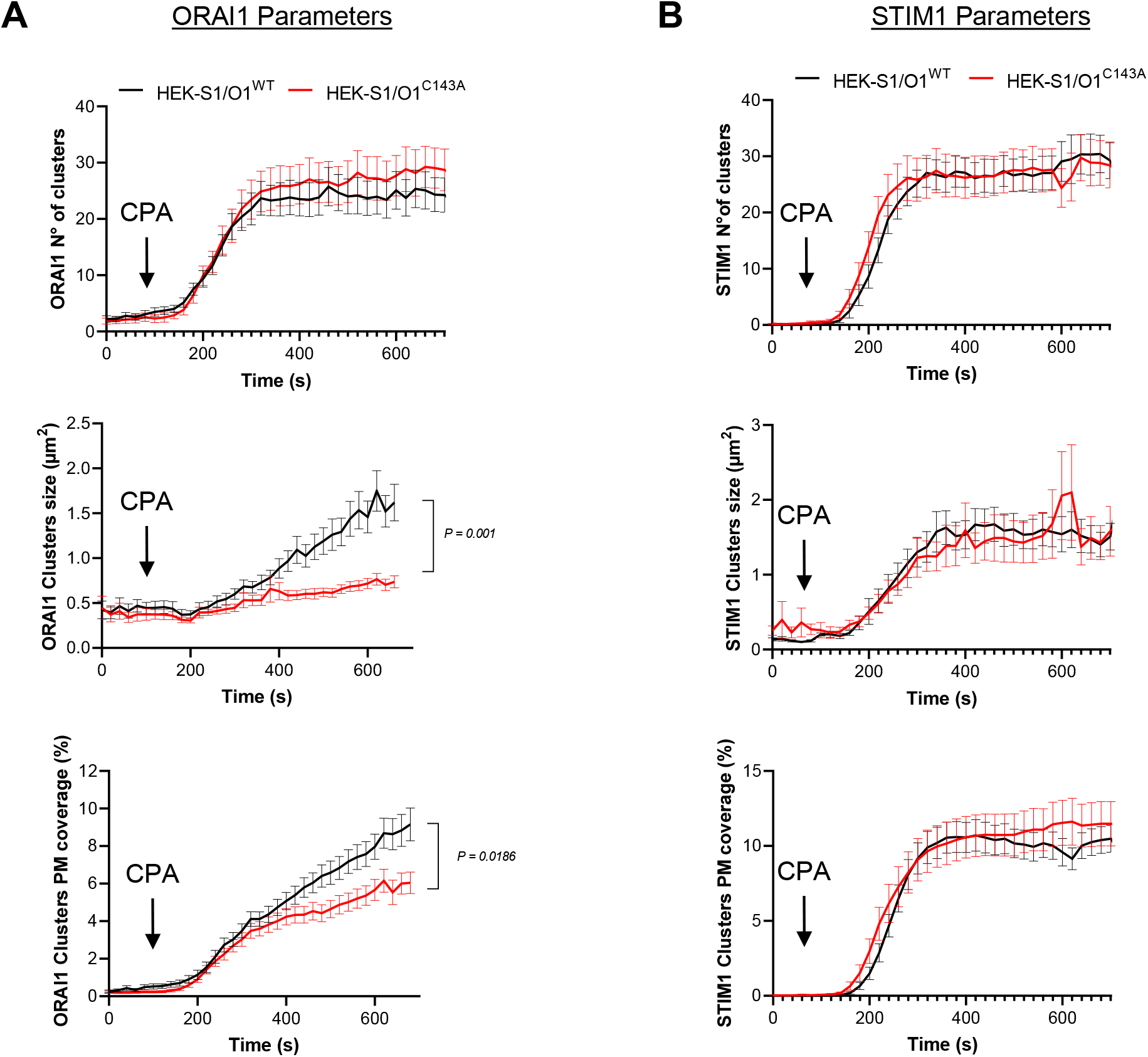
Time-course of CPA-induced changes in the number, size (μm^2^), and extent of PM covered by ORAI1-GFP (left) and mCh-STIM1 (right) clusters in WT and C1434A O1/S1 cells. Data are mean±SEM of 29 (WT) and 30 (C143A) cells from 3 independent experiments. Two-Way ANOVA fitting mixed model.

**Supplementary Fig. 4 (related to Fig. 4).**
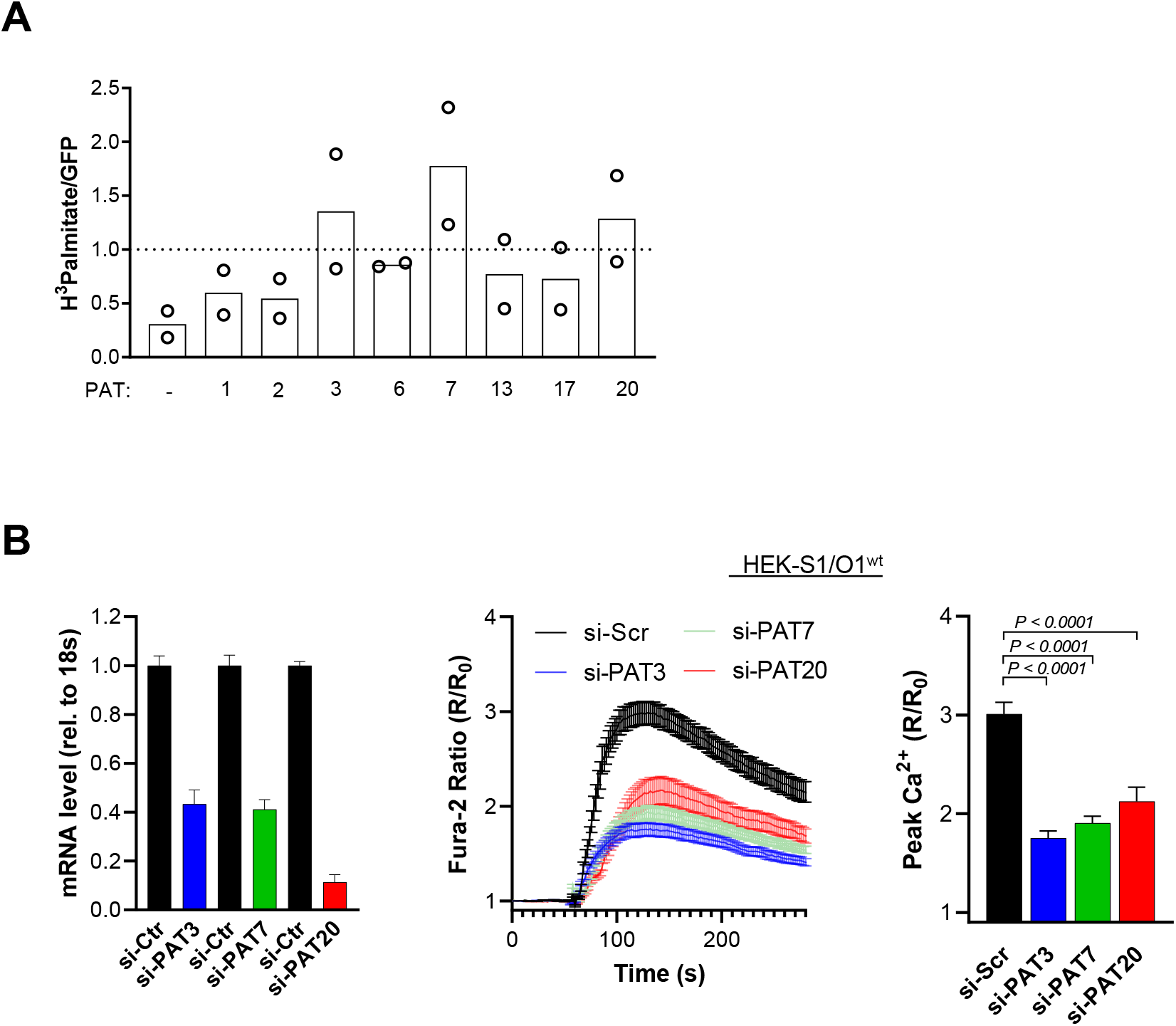
(A) _3_H-palmitate incorporation in RPE-1 cells expressing ORAI1-GFP plus the indicated PAT isoforms as in Fig. 4A, normalized for expression levels. Data are from 2 independent experiments. (B) PAT3, PAT7, and PAT20 expression levels (left), averaged SOCE responses (middle), and peak amplitude (right) in HEK-293T-S1/O1-WT cells transfected with the indicated siRNAs. Data are mean±SEM of 61-126 cells from 2 independent experiments. One way ANOVA Dunnett’s multiple comparisons test.

**Supplementary Fig. 5 (related to Fig. 5).**
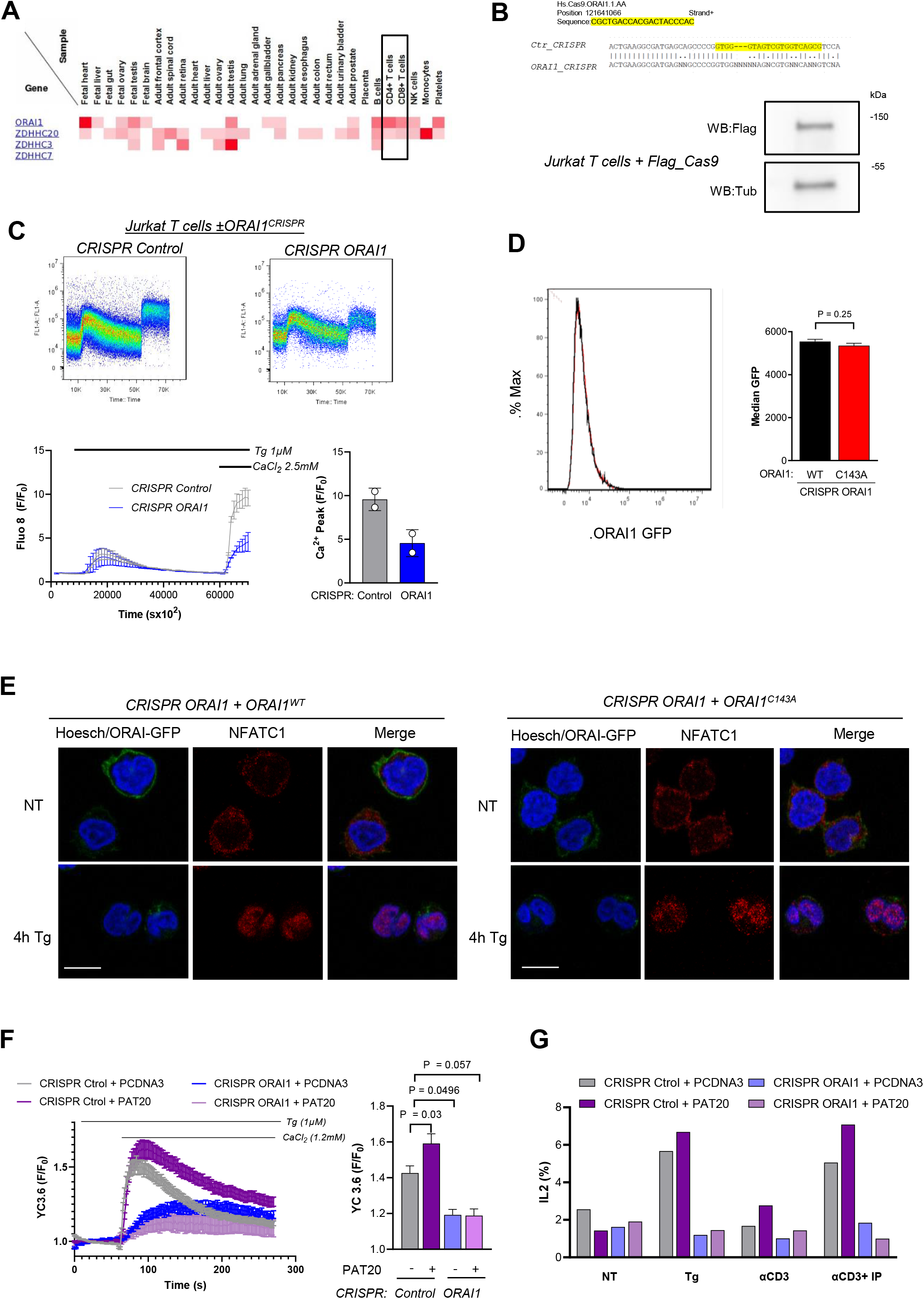
(A) ORAI1, PAT3, PAT7, and PAT20 abundance in proteomes from different tissues (from http://www.humanproteomemap.org/ consulted on Dec. 16, 2020). (B) Sequences of genomic DNA used to generate the CRISPR ORAI1 Jurkat T cell lines (top) and FLAG immunoblot of Jurkat T cells expressing FLAG-tagged Cas9 (bottom). (C) Representative flow cytometry Fluo 8 responses evoked by the Tg/Ca^2+^protocol in the indicated cells (top) and their averaged response and peak amplitude (bottom). Data are mean±SD of 2 independent experiments. (D) Fluorescence intensity profiles of CRISPR ORAI1 cells reconstituted with WT and C143 ORAI1-GFP measured by flow cytometry (N = 4). (E) Fluorescence images of CRISPR ORAI1 cells reconstituted with WT and C143 ORAI1-GFP cells blotted against NFATC1 ab treated or not with Tg to induce nuclear translocation of NFATC1 Bar = 10 µm. (F) Averaged SOCE responses (left) and peak SOCE amplitude (right) measured with YC3.6 in indicated cells expressing PAT20 or the empty vector (PCDNA3) (CRISPR Control + vector, 34 cells; CRISPR Control +PAT20, 40 cells; CRISPR ORAI1 +vector, 10 cells; CRISPR ORAI1+PAT20, 12 cells). (G) IL-2 positive cells in same cells as F treated with Tg or CD3/CD28 beads plus Ionomycin 1µM +PMA 20nM for 2h. One way ANOVA Dunnett’s multiple comparisons test.

**Supplementary Fig. 6 (related to Fig. 6).**
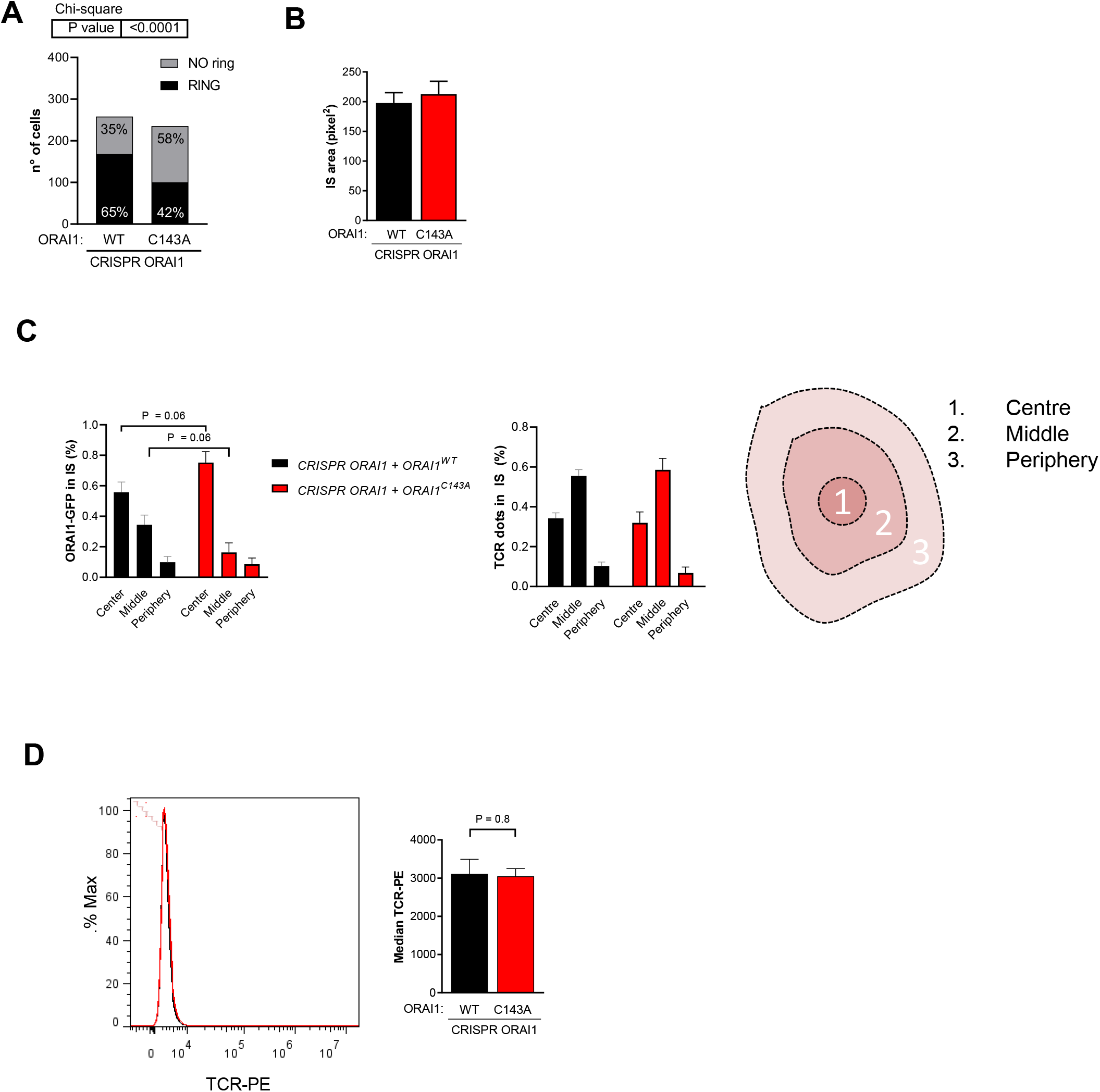
(A) Fraction of CRISPR ORAI1 cells reconstituted with WT or C143A ORAI1-GFP forming actin rings upon plating onto activating coverslips (WT = 258 cells C143A = 235 cells). (B) Averaged IS area of indicated cells forming actin rings. (C) Percentage of ORAI1 (left) and TCR (middle) signal originating from different concentric regions within the IS depicted in the sketch (right). WT = 44 cells; C143A = 27 cells. (D) TCR-PE intensity profiles of these cell lines (N = 3). Data are mean±SEM of three experiments. Two-tailed unpaired Student’s *t*-test (C) or Fisher’s exact test (A).

